# A screen for mutants deficient in coronatine-mediated suppression of root immunity identifies *Arabidopsis* SDA1 as a novel integrator of immunity and phytohormone signaling

**DOI:** 10.1101/2021.09.12.459990

**Authors:** Yi Song, Xue-Cheng Zhang, Yichun Qiu, Annika Briggs, Yves Millet, Diana Bartenstein, Catherine Mankiw, Mary L. Cerulli, Jenifer Bush, Keith L. Adams, Andrew C. Diener, Frederick M. Ausubel, Cara H. Haney

## Abstract

Despite the importance of the root immune system in the interaction with rhizosphere microbes, the majority of genetic screens for immunity regulators have been performed in leaves. A previous screen identified 27 *hsm* (<u>h</u>ormone-mediated <u>s</u>uppression of <u>M</u>AMP-triggered immunity) mutants that are impaired in jasmonic acid (JA)-mediated suppression of pattern-triggered immunity (PTI) in roots. Here we characterized 16 of the *hsm* mutants that retain JA sensitivity and are potential negative regulators of root immunity. We found that the majority of *hsm* mutants show enhanced resistance to *Fusarium*, a root fungal pathogen; however, only a subset are more resistant to a foliar pathogen. Surprisingly, 12 of 16 *hsm* mutants are also impaired in abscisic acid (ABA)-mediated suppression of PTI, suggesting a largely shared pathway between JA- and ABA-mediated immune suppression in roots. Although all *hsm* mutants are insensitive to JA-mediated suppression of root immunity, *hsm4* shows hypersensitivity to JA-mediated root growth inhibition and JA-induced gene expression. Consistently, *hsm4* is more resistant to leaf pathogens, suggesting that HSM4 is a negative regulator of both root and leaf immunity. *Hsm4* was mapped to a mutation in a conserved ARM-repeat protein homologous to yeast SDA1, which has been reported to regulate 60S ribosome biogenesis. As translational reprogramming is a critical layer of immune regulation, this work suggests that AtSDA1 is a novel negative translational regulator of immunity. Additionally, a comprehensive characterization of all 16 *hsm* mutants provides a genetic toolkit to identify novel mechanisms that regulate root immunity.

## Introduction

In plants, photosynthetically fixed carbon is secreted to the rhizosphere as root exudates, which create a nutrient- and microbe-rich niche (Sasse et al. 2018). Despite numerous studies that have systematically studied immune regulation in leaves, root immunity is largely unexplored, in part due to the inaccessibility of roots. Plant roots can respond to conserved microbe-associated molecular patterns (MAMPs) such as flagellin (Felix et al. 1999) and initiate pattern triggered immunity (PTI), including callose deposition and the expression of defense genes (Millet et al. 2010). However, emerging evidence indicates that there are differences in the regulation of leaf and root immunity. For instance, there is marked spatial distribution of MAMP responses in roots (Millet et al. 2010) and activation of a strong root immune response requires a combination of damage and MAMPs (Zhou et al. 2020). These results suggest that forward genetic screens to identify novel components of root immunity have the potential to reveal root-specific immune signaling pathways.

Activation of PTI in roots affects microbiome composition (Ma et al. 2021) and many root colonizers have evolved the ability to suppress root immunity (Yu et al. 2019; Ma et al. 2021). In contrast to suppression of PTI in leaves by pathogens, suppression of root PTI is independent of bacterial type III secretion (Millet et al. 2010). Commensal microbes utilize diverse strategies to bypass host immunity, including acidification of the rhizosphere (Yu et al. 2019), synthesis of flagellin peptide variants to block root immune receptors (Teixeira et al. 2021), and hijacking host hormone-immunity crosstalk pathways (Millet et al. 2010; Zheng et al. 2012).

Manipulation of hormone signaling is a common strategy used by microbes to suppress leaf immunity (Jones and Dangl 2006), but the mechanisms underlying hormone and immunity antagonism in roots are poorly studied. In leaves, jasmonic acid (JA) can antagonize salicylic acid- (SA) mediated defense against biotrophic pathogens, including *Pseudomonas syringae* (Loake and Grant 2007). *P. syringae* exploits this regulatory pathway by synthesizing the phytotoxin coronatine (COR), which is a structural mimic of the JA derivative JA-Isoleucine. COR suppresses SA signaling and PTI in leaves (Zheng et al. 2012). However, COR-mediated immune suppression in roots does not rely on salicylic acid-jasmonic acid antagonism (Millet et al. 2010) indicating that COR-mediated immune suppression may rely on a novel mechanism.

To identify novel regulators of root immunity, a previous genetic screen was designed to identify mutants impaired in COR-mediated suppression of PTI in roots. COR can suppress induction of a variety defense-related marker genes including *CYP71A12* (Millet et al. 2010), which is also a general non-self response (GNSR) gene essential for immunity (Maier et al. 2021). The screen made use of a transgenic PTI reporter line with the promoter of *CYP71A12* driving the *ß-glucuronidase* (*GUS*) gene (*pCYP71A12:GUS*). This screen yielded 27 *hsm* (<u>h</u>ormone-mediated <u>s</u>uppression of <u>M</u>AMP-triggered immunity) mutants that are impaired in COR-mediated suppression of *pCYP71A12:GUS* expression elicited in the root elongation zone by Flg22 (Zhang et al. 2015), which is a synthetic peptide corresponding to a conserved epitope in bacterial flagellin that activates PTI (Gómez-Gómez and Boller 2000). This screen should identify either positive regulators of JA signaling or negative regulators of PTI. A JA-insensitive mutant, *hsm1*, was previously mapped to *SGT1b* and has been shown to regulate the turnover of the COI1 receptor (Zhang et al. 2015). Another mutant *hsm13* was mapped to *FERONIA* receptor-like kinase and is a regulator of rhizosphere microbiome assembly (Song et al. 2021). These characterized *hsm* mutants confirm that this screen effectively identified regulators of hormone or immunity signaling in roots.

Of the 27 *hsm* mutants isolated, 11 were JA-insensitive, suggesting that they encode positive regulators of JA signaling (Zhang et al. 2015). Of these 11, nine are allelic to the methyl-JA receptor *COI1* and one is allelic to *MYC2* (a major transcriptional regulator of JA signaling) (Zhang et al. 2015). The remaining 16 mutants (*hsm2-hsm17*) retain JA sensitivity and are likely to be JA-responsive negative regulators of immunity in roots.

Here, we systematically characterized the growth and immunity phenotypes of the remaining 16 JA-sensitive *hsm* mutants, *hsm2-hsm17*, that we predict are negative regulators of root immunity. We found that in addition to being impaired in JA-PTI crosstalk, the majority of the *hsm* mutants are also impaired in ABA-PTI crosstalk, suggesting a shared pathway between ABA and JA antagonism with root immunity. We mapped *hsm4* to an evolutionarily conserved *SDA1* gene involved in ribosome biogenesis that may affect post translational reprogramming during immunity. Collectively, this work identified a novel negative regulator of leaf and root immunity as well as providing a genetic toolkit of additional mutants to further study the mechanisms underlying PTI-phytohormone signaling crosstalk and to identify additional novel regulators of root immunity.

## Results

### *Hsm* genes are negative regulators of root immunity

Since *hsm* mutants are predicted to be either positive regulators of JA signaling or negative regulators of root immunity, we wanted to distinguish categories of mutants by testing their hormone responsiveness. We confirmed the previous report that *hsm2-hsm17* retain sensitivity to JA-mediated root growth inhibition (Fig. 1A). In the presence of 20 µM MeJA, the parental line *pCYP71A12:GUS* exhibited approximately a 50% reduction in root length relative to mock treatment, while a mutant impaired in JA perception (*coi1-16*) only had a 22% reduction root length (Fig. 1A). We found that most *hsm* mutants showed a similar or a slightly higher degree of root growth inhibition than the parental line in the presence of MeJA. Interestingly, *hsm4* was hypersensitive to MeJA-mediated root growth inhibition with a nearly 90% reduction in root length in the presence of JA (Fig. 1A). In contrast, *hsm3* retained 61.3% root length relative to mock treatment, indicating it is less sensitive to MeJA treatment than the parental line (Fig. 1A). Collectively, these data indicate that with the exception of *hsm3*, the *hsm2-hsm17* mutants retain normal or slightly enhanced JA sensitivity, and the loss of hormone-mediated suppression of root immunity is likely due to root immune regulation rather than JA perception or signal transduction.

**Fig. 1.**
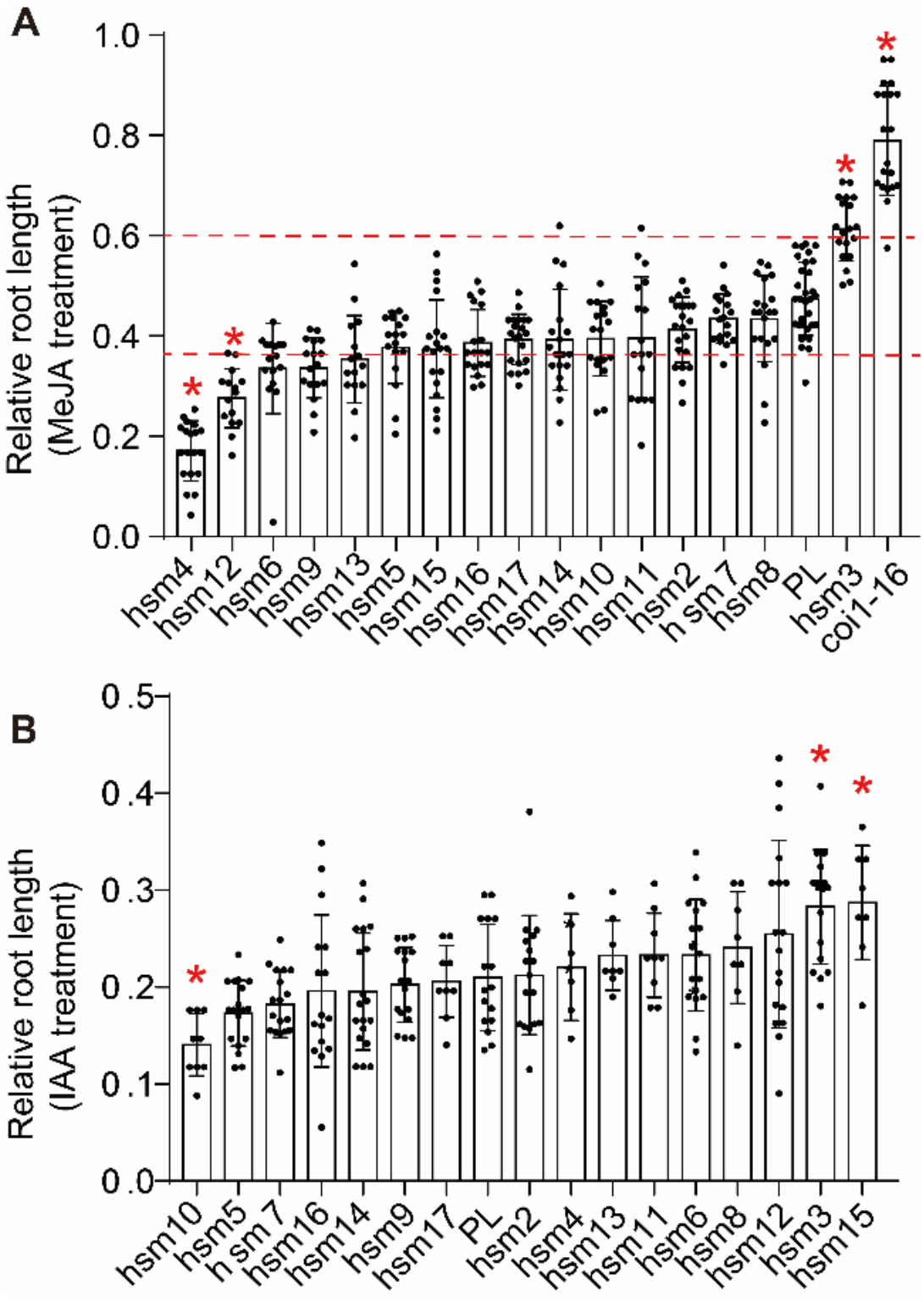
*Hsm* mutants are responsive to jasmonic acid and auxin treatment. **A**, The relative root length (treated plants/average root length of mock group) after 20 µM MeJA treatment. Data from two independent experiments, Student’s T-test followed by Benjamini-Hochberg correction was used to determine significance, all genotypes were compared with the parental line *pCYP71A12:GUS* (PL). * indicates genotypes with a greater than 25% difference (dashed red lines) from the parental line and a significant adjusted p<0.05). **B**, The relative root length after 0.5 μM IAA treatment. Seedlings were grown vertically on 1/2 strength MS plates. Root lengths were measured 8 days after germination. n>=6 for each experiment. Data from two independent experiments for 10 of 17 genotypes. Student’s T test followed by Benjamini-Hochberg correction, and for genotypes with an adjusted p value less than 0.05 were labeled with *.

Both auxin and JA are recognized by F-box family receptors that rely on the E3 ubiquitin ligase system for homeostasis (Chow and McCourt 2006; Spartz and Gray 2008), and HSM1/SGT1b is involved in the modification and turnover of both the COI1 (JA) and TIR1 (auxin) receptors (Zhang et al. 2015). We therefore tested whether the *hsm2-hsm17* mutants also affect sensitivity to auxin-mediated root growth inhibition (Fig. 1B). We found that similar to the parental line, the majority of *hsm* mutants retained 20-30% of the root length of untreated plants when grown on MS plates supplied with 0.5 μM indole acetic acid (IAA). We found two genotypes (*hsm3* and *15*) that have a significant decrease in auxin sensitivity, and only *hsm10* showed enhanced auxin sensitivity (Fig. 1B). This suggests that most of these *hsm* mutants have no or only minor deficiencies in auxin perception and that *HSM* gene products affect regulation of root immunity rather than hormone signaling. As *hsm3* shows decreased sensitivity to both JA and IAA mediated suppression of root growth, this indicates that like *HSM1/SGT1b*, it might affect a shared component of IAA and JA signaling.

To determine if the *hsm2-17* mutants exhibit aberrant immune responses in roots and leaves, we tested the susceptibility of *hsm2-hsm17* to the root fungal pathogen *Fusarium oxysporum* and the well-studied foliar pathogen *Pseudomonas syringae* pv. tomato (*Pto*) DC3000. We found that 6 *hsm* mutants (*hsm3, 6, 7, 9, 12* and *15*), are significantly more resistant to *F. oxysporum* than wild-type plants (Figs. 2A,B), suggesting that most *HSM* genes encode negative regulators of immunity in roots. We also found that whereas *hsm4, 5, 7, 13, 15* and *16* exhibit increased resistance to *Pto* DC3000 relative to wild-type plants (Fig. 2C), *hsm3, 10, 11*, and *14* are more susceptible to *Pto* DC3000. The lack of correlation between susceptibility or resistance to *F. oxysporum* and *Pto* DC3000 is consistent with previous observations that there are differences in the genetic regulation of root and shoot immunity. Collectively these results suggest that the majority of *HSM* genes encode negative regulators of root immunity.

**Fig. 2.**
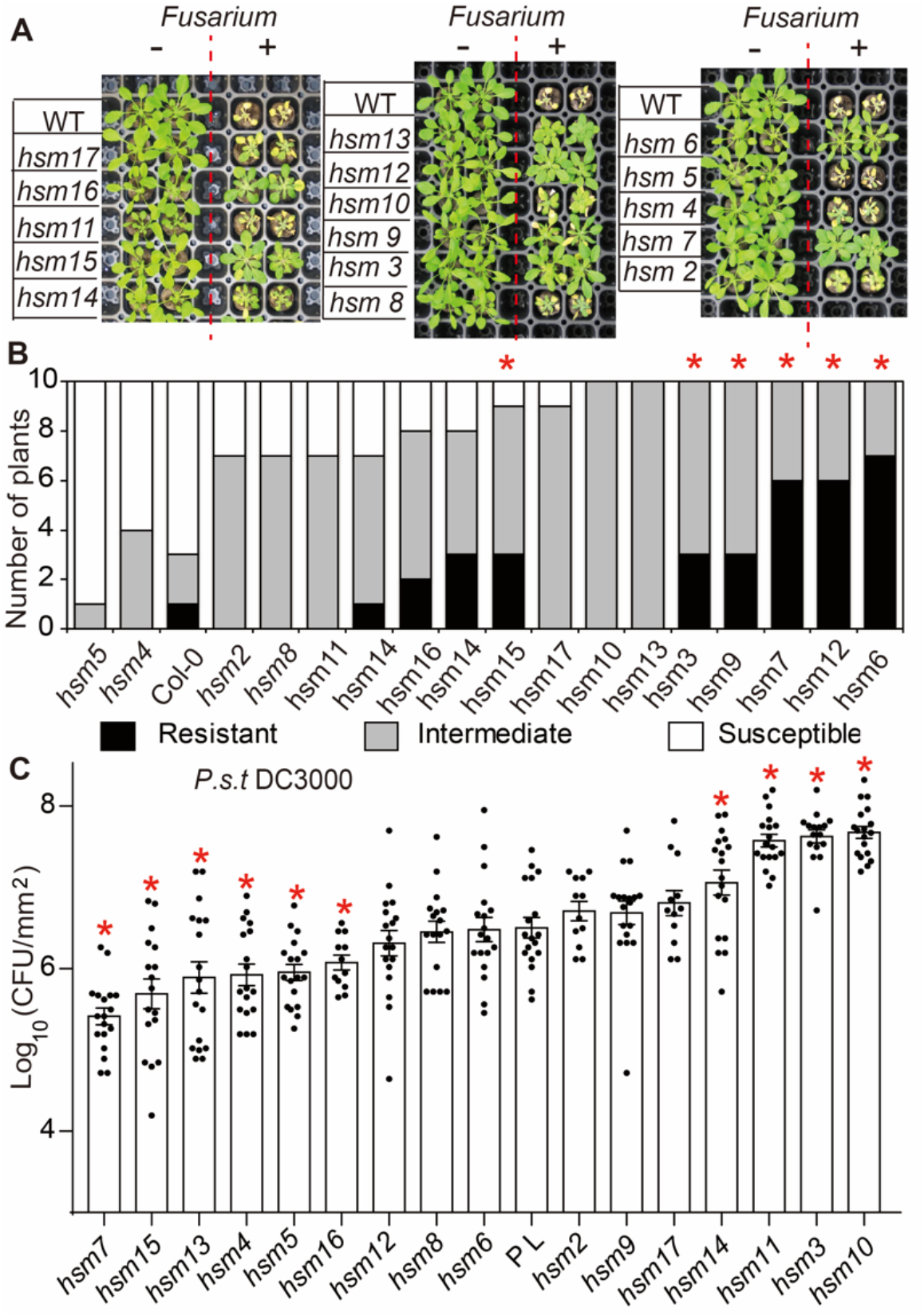
The majority of *hsm* mutants are negative regulators of root immunity. **A**, Phenotypes of *Fusarium oxysporum* infected plants. **B**, Quantification of the number of plants showing different levels of resistance to *F. oxysporum*. U test with Holm–Bonferroni Correction was used to determine the significance, and for genotypes with adjusted p < 0.05 are labeled with *. **C**, Quantification of *Pto* DC3000 growth 2 days after infiltration in the leaves of the parental line and *hsm* mutants. PL represents the parental line *pCYP71A12:GUS*. Student’s T-test compared with the parental line followed by Benjamini-Hochberg correction was used to determine the adjusted p values, *indicates mutants with p < 0.05. Dots represent CFUs from a single genotype from two (*hsm2, 16, 17)* or three (all other genotypes) independent experiments.

Mutants impaired in JA and/or immune signaling can also exhibit growth-related phenotypes (Feys et al. 1994; van Wersch et al. 2016). We found that *hsm2, hsm11*, and *hsm13/fer-8* are visibly stunted in soil, and that *hsm3* and *hsm11* have serrated leaves [Sup. Fig. 1; (Song et al. 2021)]. The severely serrated leaves of *hsm11* is similar to phenotypes observed for several loss of function mutants of microRNA genes (Laubinger et al. 2008; Lobbes et al. 2006). In contrast, we did not observe any morphological and developmental differences between *pCYP71A12:GUS* reporter line or the other *hsm* mutants and Col-0 plants. These findings indicate that a subset of the *hsm* mutants may have pleotropic effects on growth and development.

### *Hsm* mutants are largely impaired in abscisic acid-mediated immune suppression

In addition to JA, abscisic acid (ABA) can also inhibit Flg22-mediated *CYP71A1:GUS* in roots (Zhang et al. 2015). Given that ABA is a critical abiotic stress hormone, identifying mutants impaired in ABA-mediated suppression of PTI could reveal mechanisms underlying ABA-immunity crosstalk. We found that among the 16 tested *hsm* mutants, 12 mutants are also impaired in ABA-mediated suppression of Flg22-triggered expression of *pCYP71A12:GUS* (Fig. 3). The *hsm8, 9, 13* and *15* mutants were not impaired in ABA-mediated suppression of *pCYP71A12:GUS* expression, whereas *hsm12* and *hsm17* were partially impaired in ABA mediated immune suppression in roots. Because most of the *hsm* mutants are impaired in both COR- (JA-) and ABA-mediated suppression of PTI, there appears to be a substantial overlap in ABA- and JA/COR-mediated suppression of PTI in roots.

**Fig. 3.**
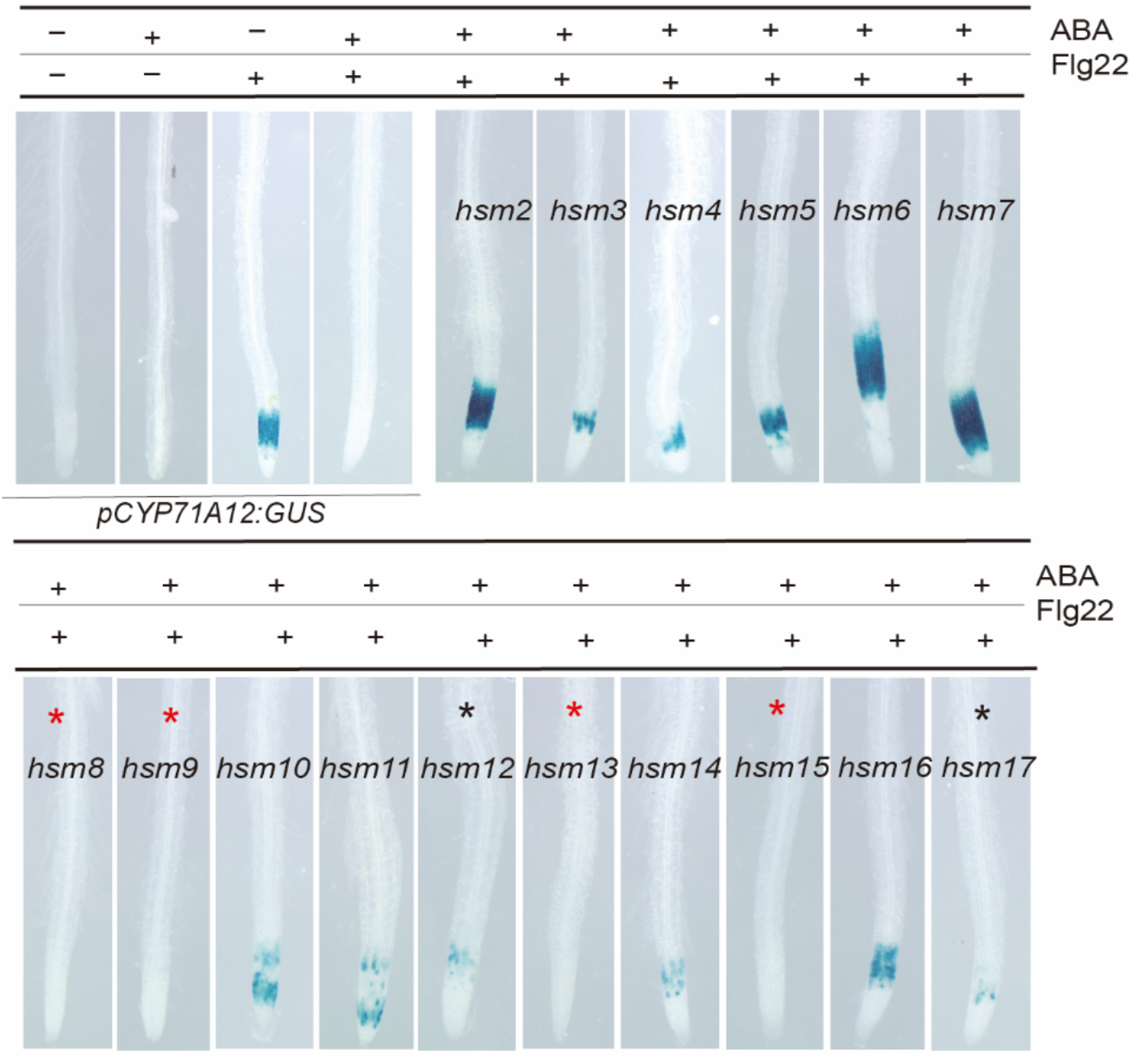
ABA-mediated suppression of root immunity is impaired in the majority of *hsm* mutants. Left top 4 panels: ABA (10 μM) strongly suppresses Flg22 (500 nM) induced *pCYP71A12:GUS* expression in the parental line. Other panels: 12 of 16 *hsm* mutants are impaired in ABA-mediated suppression of Flg22 induced *pCYP71A12:GUS* expression. Four mutants (indicated by red stars) do not affect the suppression and two mutants (indicated by black stars) partially dampen ABA-mediated suppression of PTI.

### *HSM4* encodes a putative ribosome component

We chose *hsm4* for further characterization because *hsm4* is insensitive to JA/COR-mediated suppression of root immunity but shows increased sensitivity to JA-mediated root growth inhibition, suggesting that *hsm4* might have enhanced responsiveness to MAMPs (Fig. 1A). Consistently, *hsm4* is more resistant to *Pto* DC3000, suggesting that *hsm4* encodes a negative regulator of leaf immunity (Fig. 2C). Moreover, we previously found that *hsm4* supports significantly lower levels of beneficial rhizosphere *P. fluorescens* (Song et al. 2021), suggesting that it may be a negative regulator of immunity in both shoots and roots. To test whether *hsm4* enhances sensitivity to JA-mediated transcriptional changes in roots, we treated *hsm4* and *pCYP71A12:GUS* with 1 μM COR to activate the expression of JA-responsive genes. We then measured the expression of several early (induced less than 30 minutes post treatment) JA-responsive genes (Hickman et al. 2017) in roots. We observed that the JA-responsive gene *MYC2* is more highly induced in *hsm4* relative to *pCYP71A12:GUS* (Sup. Fig. 2). Consistently, diverse JA-responsive marker genes, including *AOS, OPR3, JAZ1, JAZ3 JAZ6* and *JAZ9* were also induced to higher levels in *hsm4* roots compared with *pCYP71A12:GUS* after COR treatment (Sup. Fig. 2). These data support the conclusion that *hsm4* is hypersensitive to JA treatment at both the molecular and physiological levels.

To identify the causative mutation underlying the *hsm4* mutant phenotype, we crossed *hsm4* (in the Col-0 ecotype) to the *Arabidopsis* Landsberg ecotype. We then performed sequencing-assisted mapping in a F2 segregant population to detect chromosome regions enriched in SNPs from the Col-0 ecotype, which should be linked to the *hsm4* mutation (Methods). We identified an enrichment of SNPs from the Col-0 ecotype relative to Ler between 3.2-7.3 million bp on chromosome 1 (Fig. 4A), and we identified 7 polymorphisms in predicted coding genes present in this region in the *hsm4* genome. Two polymorphisms in AT1G13160 and AT1G15860 are predicted to cause non-synonymous amino acid substitutions in *hsm4* exons. Using publicly available transcriptional data we found that AT1G13160 is induced in response to Flg22 treatment while AT1G15860 is not (Winter et al. 2007), and so we hypothesized that a loss of AT1G13160 was more likely to underlie the *hsm4* mutant phenotype.

**Fig. 4.**
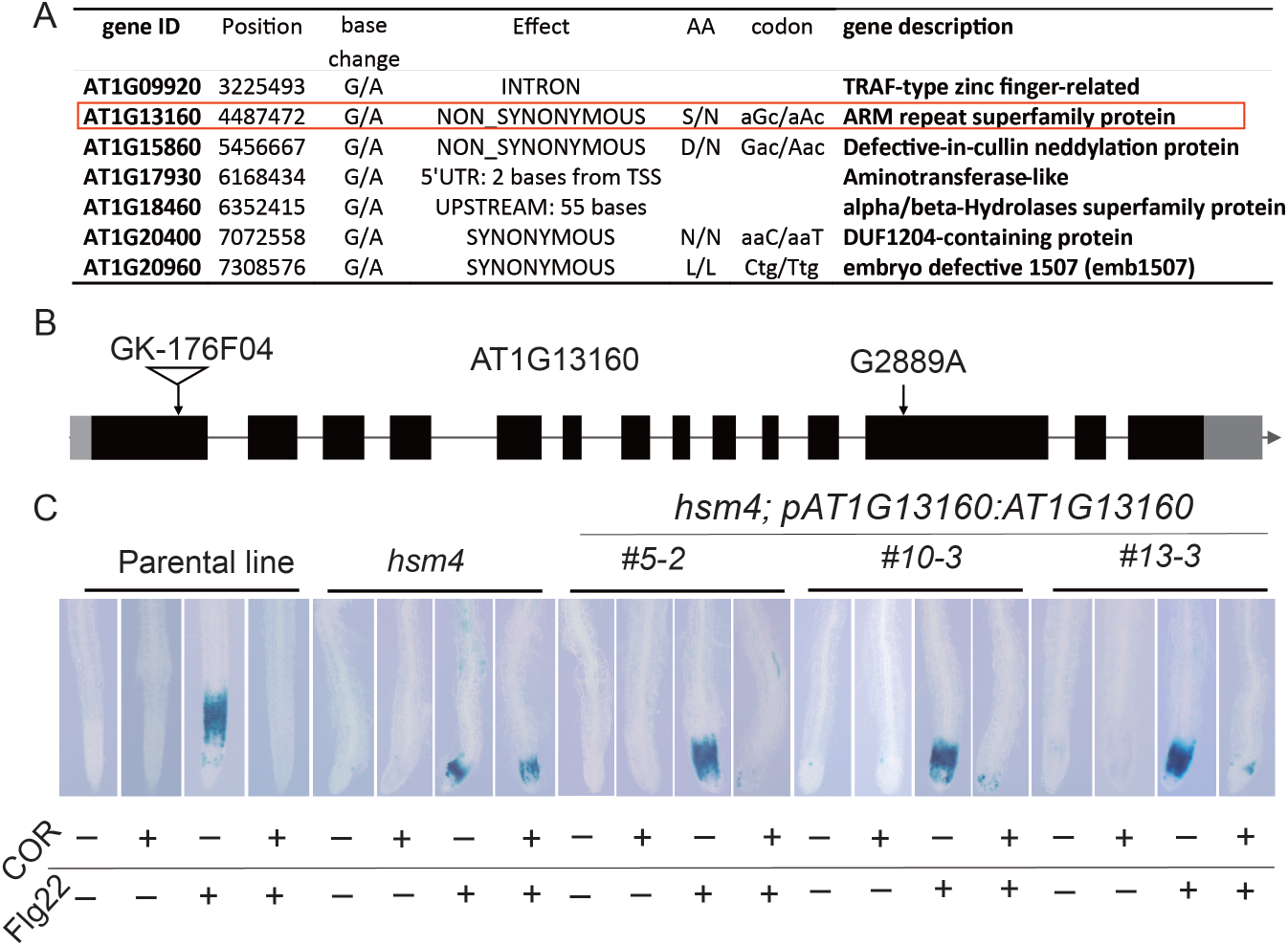
The *hsm4* mutant phenotype is due to a loss of function of in AT1G13160. **A**, Mutations in *hsm4* were mapped to a region on chromosome 1 containing 7 SNPs, and 2 of them are non-synonymous mutations. **B**, A schematic diagram shows the mutation site in *hsm4* and the available T-DNA insertion line; black regions represent exons and lines represent intron regions, light black regions represent untranslated regions (UTRs). **C**, Expression of AT1G13160 under its native promoter restored the effect of COR (2.5 μM) mediated suppression of Flg22 (500 nM) triggered *pCYP71A12:GUS* expression in 3 independent transformation lines.

To test whether loss of function of AT1G13160 underlies the *hsm4* phenotype, we expressed a full-length genomic copy including the native promoter of AT1G13160 in the *hsm4* background (Fig. 4B). Three independent complementation lines restored the effect of COR-mediated suppression of Flg22-triggered PTI in roots (Fig. 4C). We also crossed a T-DNA line with an insertion in AT1G13160 to the *hsm4* mutant. We found that the F1 progeny retained high GUS signal in roots after COR/Flg22 co-treatment (Sup. Fig. 3). In contrast, crossing *hsm4* to Col-0 showed restoration of wild-type JA-PTI crosstalk (Sup. Fig. 3), confirming that the *hsm4* mutation is recessive. These data show that the G2889A mutation in AT1G13160 (resulting a predicted S556N ammino acid substitution) underlies the *hsm4* phenotype.

Since AT1G13160 is an unstudied gene in *Arabidopsis*, we conducted a phylogenetic analysis to identify potential homologs that are functionally characterized in other organisms. We found that *HSM4* (AT1G13160) encodes a protein with 26% identity and 46% similarity (global alignment by Needleman-Wunsch algorithm) to SEVERE DEPOLYMERIZATION OF ACTIN PROTEIN 1 (SDA1) in the yeast *Saccharomyces cerevisiae*. Biochemical and genetic evidence in yeast show that SDA1 is involved in 60S ribosome biogenesis. Loss of function of *sda1* disrupts pre-RNA cleavage in response to temperature stress in yeast (Ihmels et al. 2002) and SDA1 can be co-precipitated with pre-rRNA processing and ribosome assembly related proteins (Nissan et al. 2002). Human SDA1 protein is also located in the granular component (GC) of the nucleolus, where nucleolar ribosome synthesis occurs (Babbio et al. 2004). These findings indicate that HSM4 is functionally conserved across mammals and fungi and that a defect in ribosome biogenesis may underlie the deficiency in JA-PTI crosstalk in the *hsm4* mutant.

To determine whether HSM4 shares potential functional conservation with yeast SDA1, we surveyed SDA1 homologs across eukaryotes. We found that SDA1 proteins are present in all surveyed eukaryotes and that most organisms have a single SDA1 gene in their genomes [except for *Arabidopsis* (Fig. 5A)], indicating that SDA1 is a duplication-resistant gene that has been restored to single copy status following multiple whole genome duplications in various organisms (Smet et al. 2013). Sequence alignment suggests that SDA1 homologs among eukaryotes have several conserved domains (Sup. Fig. 4).

**Fig. 5.**
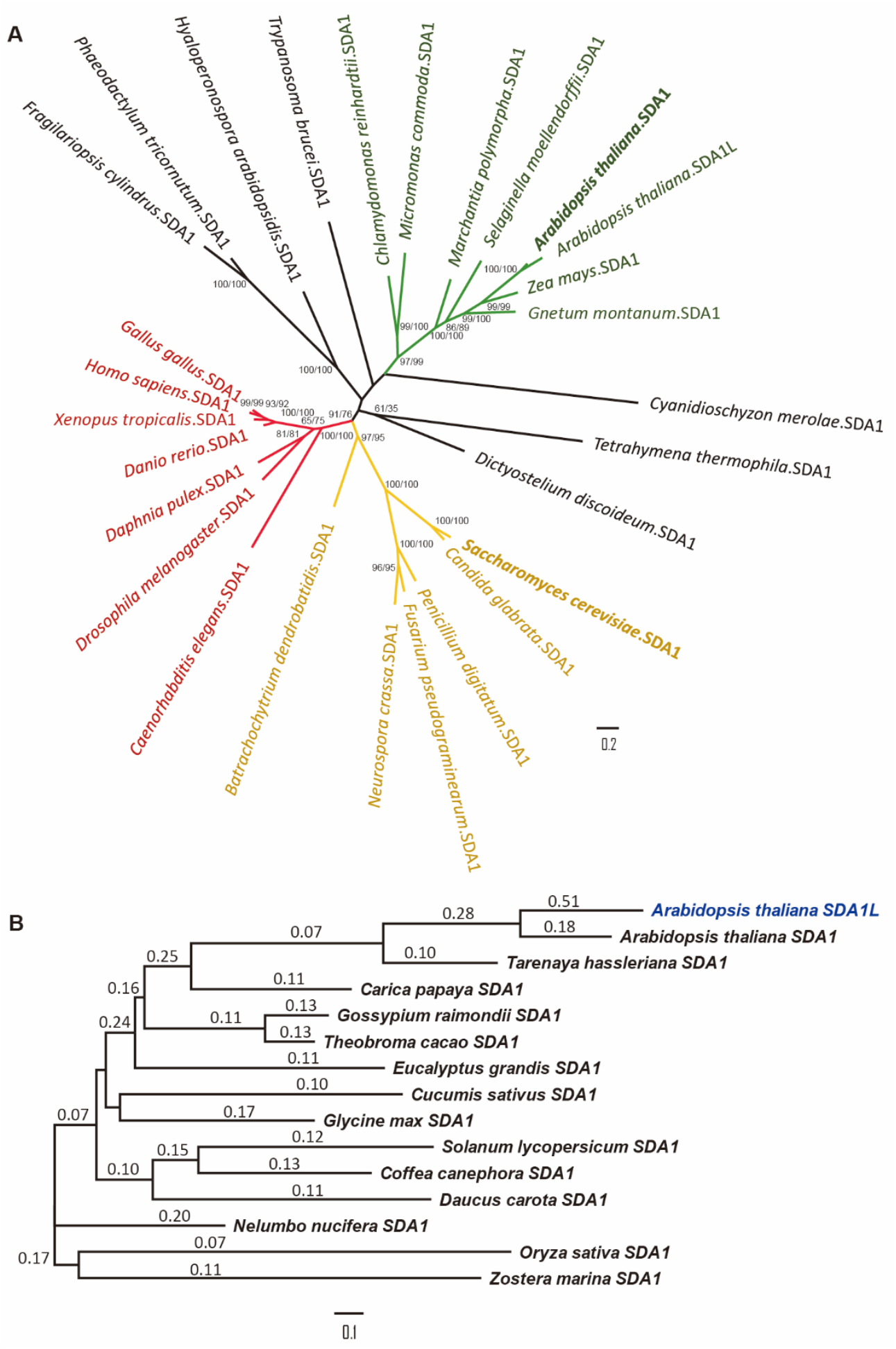
Phylogeny of the eukaryotic SDA1 gene family. **A**, Numbers closest to nodes are bootstrap (%) values calculated by SH-aLRT and ultrafast bootstrapping methods, respectively. Green indicates the plant lineages including *HSM4* in bold, the gene of interest in this study. Red indicates the animal lineages. Yellow indicates the fungal lineages including the yeast SDA1 in bold, the homolog with reported functions. Black indicates other protists. SDA1 homologs are present as single-copy genes in most of the surveyed eukaryotes. The scale bar indicates the substitution rate per site. **B**, Sequence evolution rates of genes homologous to SDA1 in angiosperms. The Ka/Ks values are labelled above the branches. The scale bar indicates the substitution rate per codon. The divergent paralog in *Arabidopsis thaliana* is shown in blue. All SDA1 genes are under purifying selection with a low Ka/Ks value.

We reasoned that if SDA1’s function is conserved in different organisms, it should be under strong purifying and/or negative selection. We therefore analyzed the Ka/Ks ratios (Yang and Bielawski 2000), which refers to the ratio of the number of nonsynonymous substitutions per non-synonymous site (Ka) to the number of synonymous substitutions per synonymous site (Ks). Low Ka/Ks ratios indicate the protein is not tolerant to most non-synonymous mutations and that the protein is under purifying selection (evolutionary conserved). Significantly, the Ka/Ks ratios of SDA1 from most organisms are very low (less than 0.2) and it is only 0.18 for AtSDA1 (Fig. 5B), indicating that the *SDA1* gene family has been under strong purifying selection during the evolutionary history of angiosperms. Thus, AtSDA1 likely functions in ribosome biogenesis similar to its yeast homolog.

Although SDA1 is a single copy gene in most organisms, we found that *HSM4* (*AtSDA1*) has a paralog in the *Arabidopsis* genome, AT4G31520 (SDA1-like, or SDA1L), which arose relatively recently after the divergence between the sister families, *Brassicaceae* and *Cleomaceae* (Fig. 5 A, B). In contrast to the *SDA1* genes from most organisms that we found have low Ka/Ks ratios (suggesting evolutionary conservation), *AtSDA1L* has a significantly higher Ka/Ks ratio (0.51), suggesting relaxation of purifying selection and putative functional divergence (Fig. 5B). Whether AtSDA1L has gained a novel function in *Arabidopsis* remains to be elucidated, but in any case, the *Arabidopsis* SDA1 and SDA1L proteins do not appear to be functionally redundant.

## Discussion

Genetic and molecular studies during the past several decades have substantially advanced our understanding of plant immune pathways; however, most studies have focused on aboveground tissues. Roots are exposed to the nutrient and microbe rich soil environment, but the conserved and distinct immune responses between roots and shoots are poorly studied. The *hsm* screen is relatively unique in that it focused on identifying novel regulators of hormone-immunity crosstalk in roots (Zhang et al. 2015). Root-specific negative regulators of immunity might have implications for immune responses to root pathogens or commensals. Our work systematically examined the growth- and immunity-related phenotypes in *hsm* mutants and found that many HSMs are required for the immune response to leaf or root pathogens. We identified AtSDA1 as a novel negative regulator of root and leaf immunity. Collectively, this work describes a genetic toolkit for further exploration of immune regulation in roots.

In eukaryotes there are two parts of the ribosome: the 40S subunit is responsible for decoding the mRNA and the 60S subunit catalyzes the peptidyl transferase reaction (Browning and Bailey-Serres 2015). Both 60S and 40S pre-ribosome assembly requires diverse non-ribosomal proteins to form a protein-RNA complex in the nucleolus, which is exported to the cytoplasm for maturation (Nissan et al. 2002). In yeast, tagged SDA1 protein was co-purified together with the 60S pre-ribosomal particles, suggesting it is a regulator of 60S ribosome biogenesis. In this work, we found that *HSM4* encodes the *Arabidopsis* SDA1 ortholog. Emerging evidence suggests that translational reprogramming is essential for immune regulation in plants. For example, ribosome footprinting analysis found that at the gene level, changes in transcription and translation rates are poorly correlated following treatment with the immune elicitor elf18, suggesting that regulation of mRNA translation is a critical layer of immune regulation (Xu et al. 2017). Since ribosome function is essential for the control of translational efficiency (Merchante et al. 2017), *HSM4* may play an important role in gating the translation of PTI-related genes in roots. However, this conclusion raises the question of how a general translation apparatus component can specifically affect innate immunity-related phenotypes without general growth and developmental defects.

Previous genetic evidence suggests that COR/JA-mediated root immunity suppression is dependent on canonical JA perception and signaling components Table 1). Mutants deficient in JA perception (*coi1-1*) and the signaling component *MYC2* (*jin1-7*) are impaired in COR/JA mediated suppression of *CYP71A12pro:GUS* expression in roots (Millet et al. 2010). MYC2 is also a component of the ABA signaling pathway and its expression is induced by ABA and drought (Kazan and Manners 2013). Consistently, 12 of 16 tested *hsm* mutants are impaired in both ABA- and COR-mediated *pCYP71A12:GUS* activation, suggesting that most *hsm* mutants act downstream of MYC2 and that there is also a substantial overlap between ABA- and JA-mediated PTI suppression. Moreover, MYC2 is not only a master regulator of JA/ABA signaling (Dombrecht et al. 2007), but also mediates crosstalk with other signaling pathways such as gibberellic acid-, ethylene-, and light-mediated signaling (Hong et al. 2012; Lorenzo et al. 2004; Yadav et al. 2005). This explains why multiple phytohormones can suppress PTI in roots (Zhang et al. 2015) and indicates that MYC2 is a signaling hub at the intersection of crosstalk between multiple hormonal signals and root immunity. A recent study also supports the crucial role of MYC2 in integrating shoot light stress and reshaping the rhizosphere microbiome (Hou et al. 2021). Further investigating the function of HSM proteins will presumably provide additional insights into pathways required for shaping interactions with root-associated pathogens and commensals.

**Table 1.**
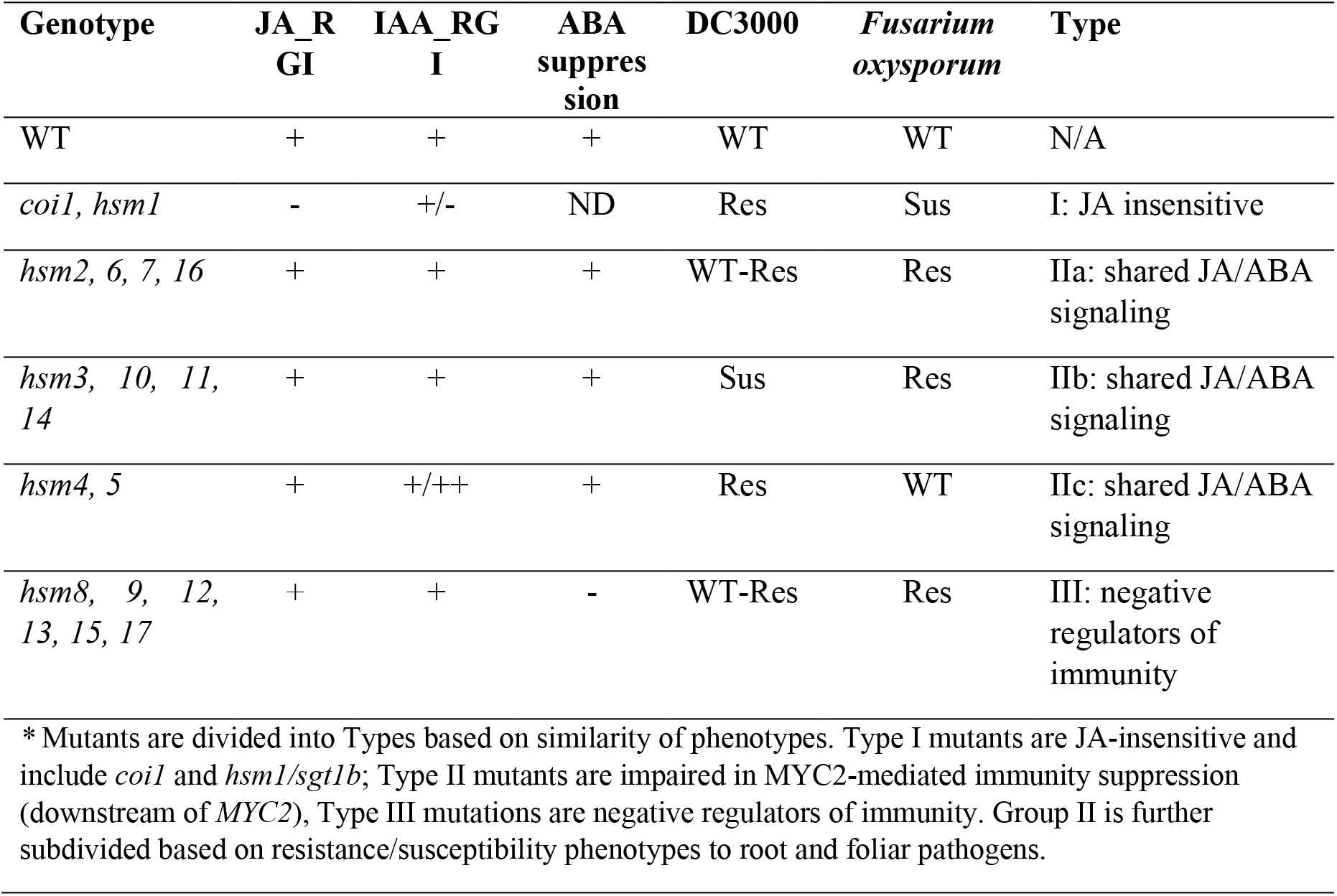
A summary of the major phenotypes of *hsm* mutants*.

We postulated that *hsm* mutants either positively affect JA signaling or negatively regulate immunity. Our data support the existence of three major categories of *hsm* mutants (Table 1): Type I mutants are JA-insensitive mutants impaired in JA induced MYC2 activation (upstream of *MYC2* and would include *coi1* and *hsm1/sgt1b*). Type II mutants are insensitive to both COR- and ABA-mediate-mediated immunity suppression. They are most likely root immune regulators downstream of *MYC2* regulated by both ABA and JA signaling. Type III mutations are negative regulators of PTI and thus may enhance sensitivity to Flg22. Since 12 *hsm* mutants show defects in both ABA and JA mediated PTI suppression, it seems likely that these mutants are impaired in a shared ABA/JA-mediated immunity suppression pathway and are Type II *hsm* mutants (impaired downstream of *MYC2*). The remaining 5 *hsm* mutants, including the previously described *hsm13* mutant, which is due to a loss of function of *FERONIA* receptor-like kinase, are still responsive to ABA mediated PTI suppression and are JA sensitive, suggesting that they are negative regulators of immunity in roots (Type III). Thus, our work on *hsm* mutants provides new phenotypic and mechanistic insights into the regulation of root immunity and the role of hormone signaling pathways in that process.

## Materials and Methods

### Plant material, growth conditions and hormone treatments

For all *hsm* mutants, F2 segregants displaying the *hsm* phenotype that did not show phenotypic segregation in the F3 generation were considered to be homozygous mutants and used for further analyses. Seeds were treated with 70% ethanol (2 min) and 10% bleach (2 min), and then washed three times in sterile water. Sterilized seeds were stored at 4°C for 2 days before planting. Plants were grown under 12 h light/12 h dark (75-90 μE) and a 23/20°C day/night temperature. For root growth inhibition assays, seeds were grown on ½ x Murashige and Skoog (MS) plates with 1% sucrose. Hormones were added to the media to a final concentration of 20 μM MeJA (Sigma) or 0.5 μM IAA (VWR). Seedlings were grown on the plates vertically in the growth room, and the root length was measured at 7 days after sowing. The relative root inhibition ratio was calculated as the length of the hormone-treated group relative to that of the mock-treated control group.

For root GUS signal detection, assays were performed as described (Millet et al. 2010). Seeds were grown in 96-well plates with 120 μL 1X liquid MS medium (pH 5.7) and 0.5% sucrose (3-5 seeds per plate). Plates were incubated on shelves at 16 hours light per day for 8 days. On day 8, the media was replaced with fresh media. Hormones were added to the indicated final concentrations, and Flg22 was added to a final concentration of 500 nM. After 6 hours of incubation with hormones and Flg22, the media was replaced with 150-200 μL GUS staining solution (50 mM pH 7 sodium phosphate buffer, 10 mM EDTA, 0.5 mM potassium ferricyanide, 0.5 mM potassium ferrocyanide, 0.5 mM x-gluc, 0.01% triton X-100), and incubated for 3 hours at 37°C.

### Pathogen infection assays

*Fusarium oxysporum* infections were performed as described in (Diener and Ausubel 2005). For *Pto* DC3000 infection assays, bacteria were grown in LB media overnight and cells were spun down and washed with 10 mM MgSO_4_. Leaves from four-week old plants were used for the assay and infiltrated with *Pto* DC3000 at a concentration of OD_600_ = 0.0002 (approximately 1000 CFU/cm^2^ leaf tissue). Plants were kept covered with a clear dome for 2 days and kept at 12h light/12 hour dark and 7-mm leaf punches were used for bacterial quantification. Leaf disks were homogenized, serially diluted and plated on rectangle plates containing LB with 50 µg/ml rifampicin for CFU calculation.

### Genome resequencing-assisted mapping and cloning of *HSM4*

The *HSM4* gene was cloned using a genome resequencing–assisted mapping method. Briefly, *hsm4* was outcrossed to Ler to yield an F1 hybrid, followed by self-fertilization to produce F2 segregants. A total of 71 F2 segregants were selected on the basis of retained GUS reporter activity in roots following COR and Flg22 co-treatment. Genomic DNA from these 71 segregants was extracted, pooled and subjected to library preparation for Illumina-based next-generation sequencing. Mapping and analysis of next the sequencing data was performed as described (Zhang et al. 2014). For genetic complementation tests, the full length genomic region covering AT1G13160 and the native promoter was cloned into the plant expression vector pPLV27 using a ligation-independent strategy (De Rybel et al. 2011). Constructs were transformed into *Agrobacterium tumefaciens* GV3101 and plants were transformed using the floral dip method. Transformed seeds were selected on 50 μg/ml kanamycin plates, and three independent homozygous T3 lines (#5-2, #13-3 and #10-3) were used for phenotyping.

### qPCR analysis

Seedlings were grown in 12-well plates in 1X liquid MS medium (pH 5.7) supplemented with 0.5% sucrose and 0.5 g/L MES hydrate, and plates were kept in 16 hours light chamber with a light intensity of 100 μE at 22 °C. The media was changed at day 8 and root samples were harvested 30 mins after 1 μM COR treatment on day 10. The gene expression levels were quantified using qRT-PCR and normalized to internal control gene *EF1α*.

### Phylogenetic and selection analyses

Amino acid sequences and nucleotide sequences of AT1G13160 and its paralog AT4G31520 were obtained from the *Arabidopsis* information resource (TAIR10; arabidopsis.org). The sequences of orthologous genes in several plant lineages were obtained in PLAZA 4.0 [https://bioinformatics.psb.ugent.be/plaza/; (Van Bel et al. 2018)] through the gene family database and confirmed by reciprocal best BLAST hits. The amino acid sequence of the yeast homolog, SDA1 from *Saccharomyces cerevisiae*, was obtained from Uniprot [https://www.uniprot.org/uniprot/P53313; (The UniProt Consortium 2017)]. The homologs from several animals, fungi and protists were identified by Inparanoid [http://inparanoid.sbc.su.se/cgi-bin/gene_search.cgi?id=P53313; (Sonnhammer and Ostlund 2015)], and the absence of additional homologs was confirmed through top BLAST hits. Homologous amino acid sequences of these taxa were also obtained from Uniprot. Each homolog was globally aligned to the *Arabidopsis* SDA1 sequence with Needleman-Wunsch algorithm to determine the sequence identity, similarity and indels (Needleman and Wunsch 1970).

The amino acid sequences of selected eukaryotic SDA1 homologs were aligned by MUSCLE with the default settings (Edgar 2004). A maximum likelihood tree was generated by the web-based IQ-TREE 1.6.7 (Nguyen et al. 2015; Trifinopoulos et al. 2016). The implemented ModelFinder determined LG to be the best substitution model (Kalyaanamoorthy et al. 2017), and 1000 replicates of ultrafast bootstrap were applied to estimate the support for reconstructed branches (Hoang et al. 2018), as well as 1000 replicates of the SH-like approximate likelihood ratio test (SH-aLRT) (Guindon et al. 2010). The tree topology and branch supports were reciprocally compared with and supported by another maximum likelihood tree generated using RAxML v. 8.1.2 (Stamatakis 2014).

To generate the codon alignment, a separate amino acid alignment of selected angiosperm SDA1 homologs was generated and used as a reference. This codon alignment was used to generate a maximum likelihood tree with GTRGAMMA as the substitution model. Several analyses were performed using Codeml in the PAML package (Yang 2007) with the codon alignment and the codon tree as input. A free-ratio test was applied to estimate the branch-wise Ka/Ks ratios for each of the phylogenetic tree branches. Additional models were applied to compare the evolutionary rates between the paralogs in *A. thaliana* and between the *A. thaliana* genes and the orthologs, and likelihood ratio tests were used to suggest the presence or absence of significant difference in the selection strength among members of the *SDA1* gene family.

## Statement of author contributions

X-C. Z., Y. S., Y.M., C.H. and F. M. A. designed experiments; X.Z., Y.S., A.B., Y.M., D.B., C.M., M.C., A.D. and J.B. performed experiments; X.Z., Y.S., Y.M., Y.Q., A.D. and C.H. analyzed data, Y.Q. and K.A. performed phylogenetic analysis; Y.S. and C.H. wrote the manuscript with input from all.

