## Supplementary Data for "A screen for mutants deficient in coronatine-mediated suppression of root immunity identifies *Arabidopsis* SDA1 as a novel integrator of immunity and phytohormone signaling"

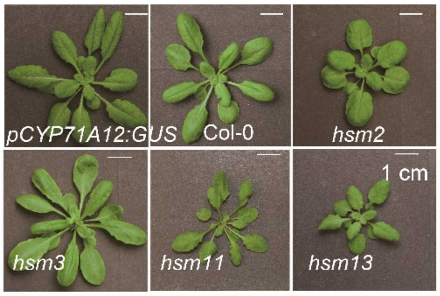


**Supplementary fig. 1.** Four *hsm* mutants show visible morphological changes compared with the parental line *pCYP71A12:GUS* and Col-0. Images are from 4-week old soil-grown plants. Phenotypes were consistent among all plants and were not phenotypically segregating.

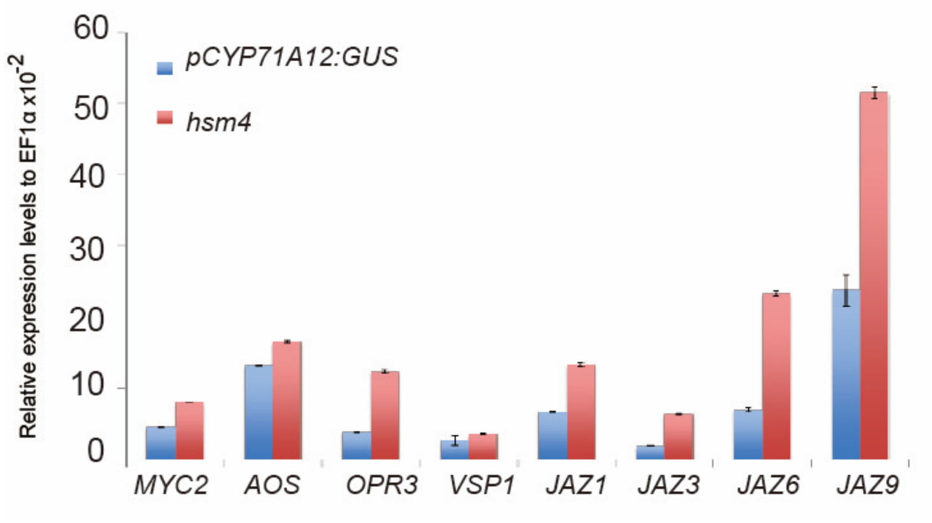


**Supplementary fig. 2.** JA-responsive genes have higher expression after COR treatment in *hsm4* relative to wild-type plants. Root samples were harvested 30 mins after 1 μM COR treatment and expression levels were quantified using qRT-PCR and normalized to internal control gene *EF1α*.

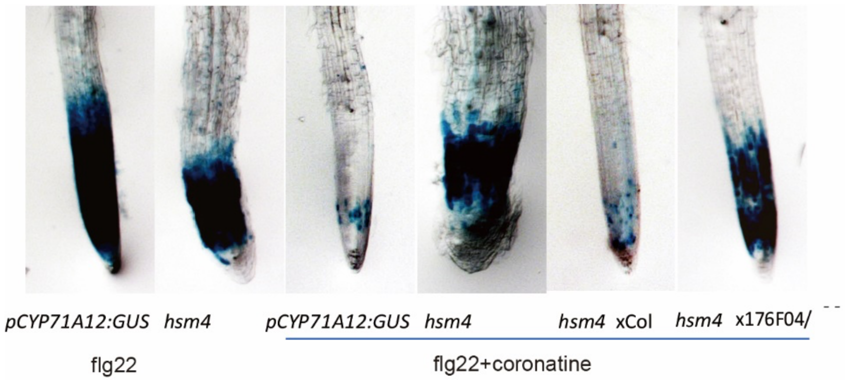


**Supplementary fig. 3. A T-DNA insertion line (GK-176F04) in AT1G13160 failed to complement the *hsm4* mutant.** *Hsm4* was crossed to a T-DNA insertion in AT1G13160 or wild-type Col-0, and F1 progeny were scored for COR-mediated suppression of *pCYP71A12:GUS* expression. While an *hsm4* × Col-0 cross restored COR-mediated suppression of root immunity, the AT1G13160 T-DNA insertion did not suggesting that the *hsm4* mutant has a loss of function of AT1G13160.

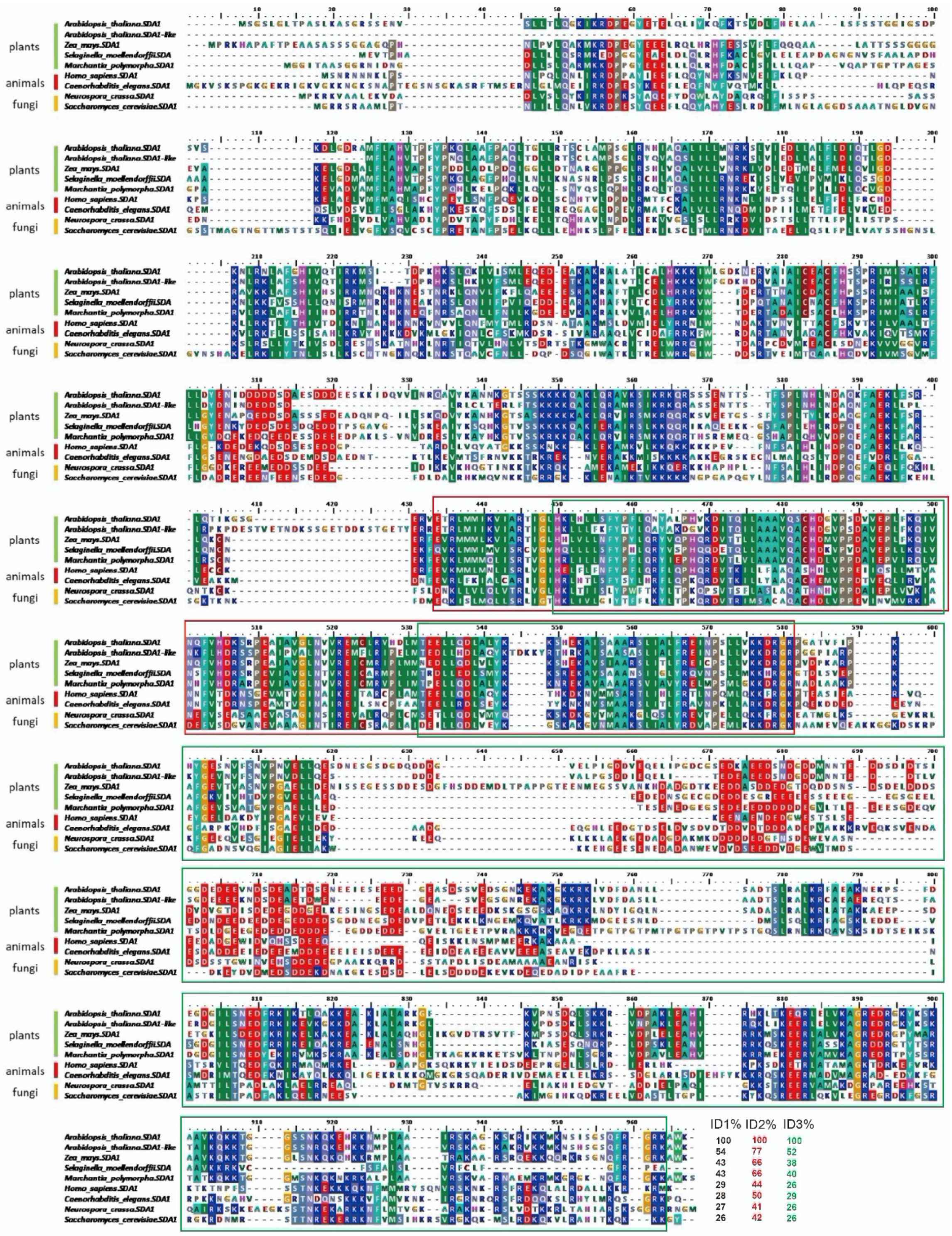


**Supplementary Fig. 4.** Alignment of selected SDA1 protein sequences. Amino acids with over 50% identity and similarity score based on the BLOSUM62 matrix are highlighted. Percent identity (ID%) 1, 2 and 3 represent full-length global alignment, Ath 352E-494R conserved and no gap region (black box), and SDA1-domain (449M-801K), respectively. The amino acids numbers of different domains are based on their positions in AtSDA1.
